## Supplementary Materials for "Plk1 overexpression suppresses tumor development by inducing chromosomal instability"

### Supplementary Figures

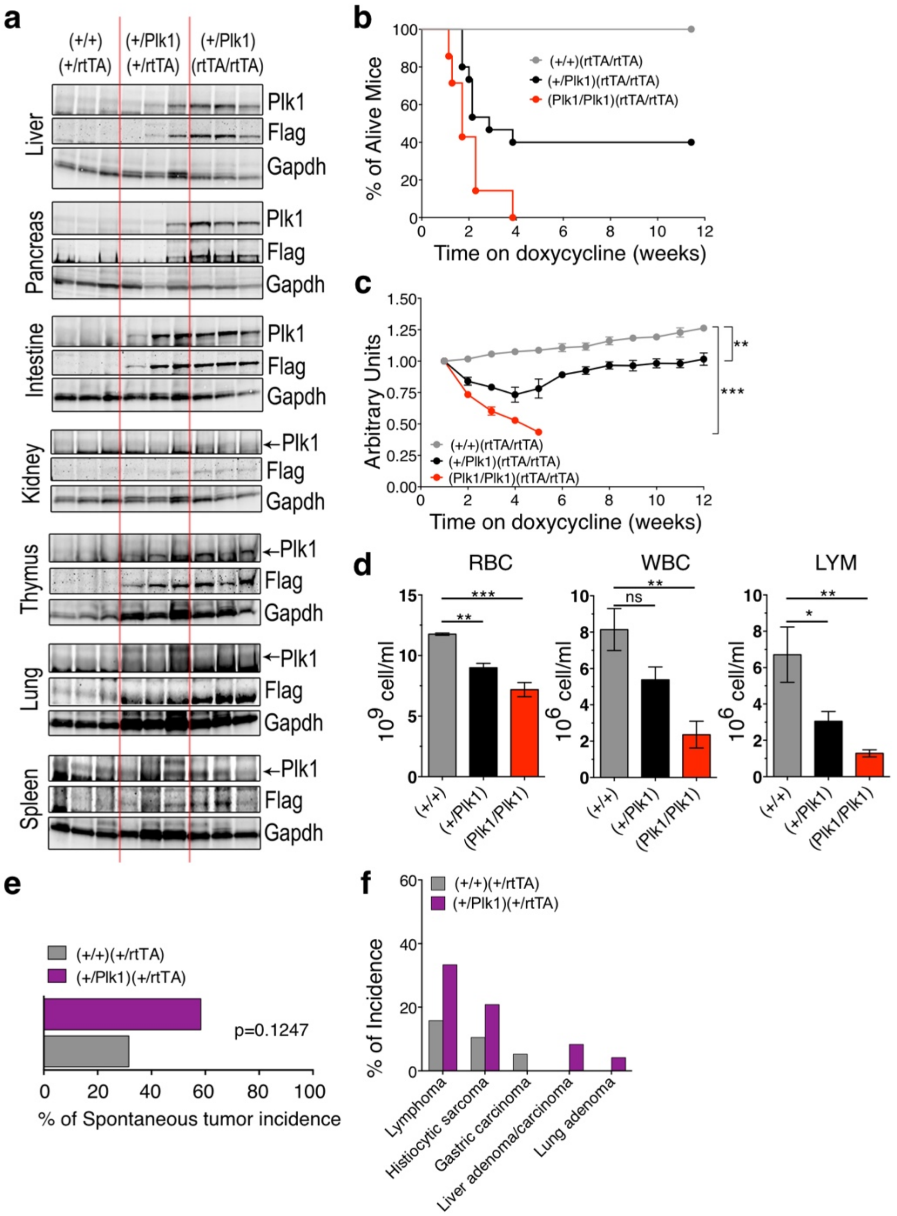


**Supplementary Fig. 1**. Inducible expression of Plk1 in mice and tumors generated. **a** Immunodetection of Plk1 in tissues from mice with the indicated genotypes fed with Dox food for 12 days. Gapdh was used as loading control. **b** Ageing analysis of Plk1 overexpressing high dosage of Plk1 (Plk1/Plk1)(rtTA/rtTA) [red line] (n=7) and (+/Plk1)(rtTA/rtTA) [black line] (n=15) compared to non-expressing mice (+/+)(rtTA/rtTA) [grey line] (n=5). **c** Normalized weigh measurements of mice depicted in panel a; (mean ± sem). ** p<0.01; *** p < 0.001; one-way ANOVA Bonferroni test. **d** Peripheral blood populations after 5 days on Dox (Plk1/Plk1)(rtTA/rtTA) (n=3), (+/Plk1)(rtTA/rtTA) (n=5) compared to non-expressing mice (+/+)(rtTA/rtTA) (n=3). Red blood cells (RBC), white blood cells (WBC) and lymphocytes (LYM). * p<0.05; ** p<0.01; *** p<0.001 one-way ANOVA Dunnett's multiple comparison test **e** Spontaneous tumor incidence in ageing mice from Fig 1e. (+/+)(+/rtTA), 19 mice; (+/Plk1)(+/rtTA), 24 mice. Distribution of tumors in ageing animals with higher levels of Plk1 (purple columns) compared to control animals (grey columns) from Fig 1e. In **e,f** differences are not significant; Fisher exact test.

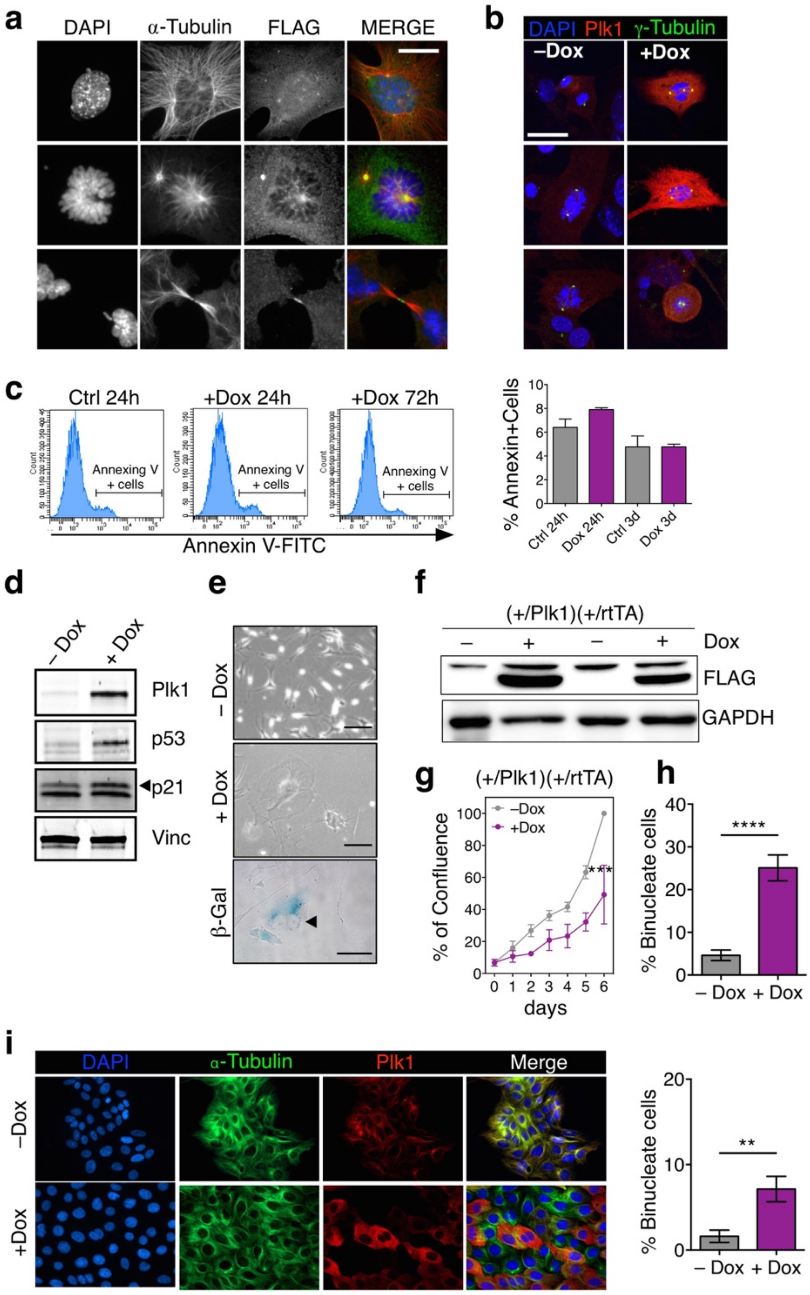


**Supplementary Fig. 2**. Inducible expression of Plk1 in MEFs and MCF10A and associated phenotypes. **a** Immunofluorescence of (+/Plk1);(+/rtTA) MEFs treated with Dox for 24h. The FLAG tag is detected at the centrosomes, spindle poles, kinetochores and the cytokinesis bridge in agreement with the known localization of Plk1 in dividing cells. **b** Immunofluorescence of MEFs in mitosis treated with Dox for 24h or untreated (–Dox). **a,b** Scale bar, 10 μm. **c** Flow cytometry analysis of cells stained for Annexin V-FITC in MEFs overexpressing Plk1 at 24h and 72h after Dox addition and control cells. Right histogram represents the percentage of Annexin V positive cells. There are no statistical significant differences in control cells versus Dox treated cells. **d** Immunodetection of the indicated antigens in (Plk1/Plk1);(+/rtTA) MEFs after 48h on Dox. Vinculin was used as a loading control. **e** Representative images of (Plk1/Plk1);(+/rtTA) cells stained for senescence-associated β-galactosidase (arrowhead) after 7 days in the presence of Dox. **f** Expression of FLAG in Hras+E1A transformed MEFs in two different clones after 24h on Dox (+) or in normal media (–). **g** Quantification of confluence in transformed cultures in the absence (–Dox) or presence (+Dox) of Dox for 6 days. ***, $p<0.001$; Two-way ANOVA. **h** Quantification of binucleated cells in Hras + E1A transformed MEFs untreated (–Dox) or treated with Dox for 48h (+Dox). ****, $p<0.0001$ (n=2 replicates); One-way ANOVA. **i** Immunofluorescence in MCF10A-rtTA cells untreated (–Dox) or treated with Dox for 48h (+Dox) to express Plk1 and quantification of binucleated cells. **h** and **i** >250 cells from at least 20 random microscope fields, are quantified.

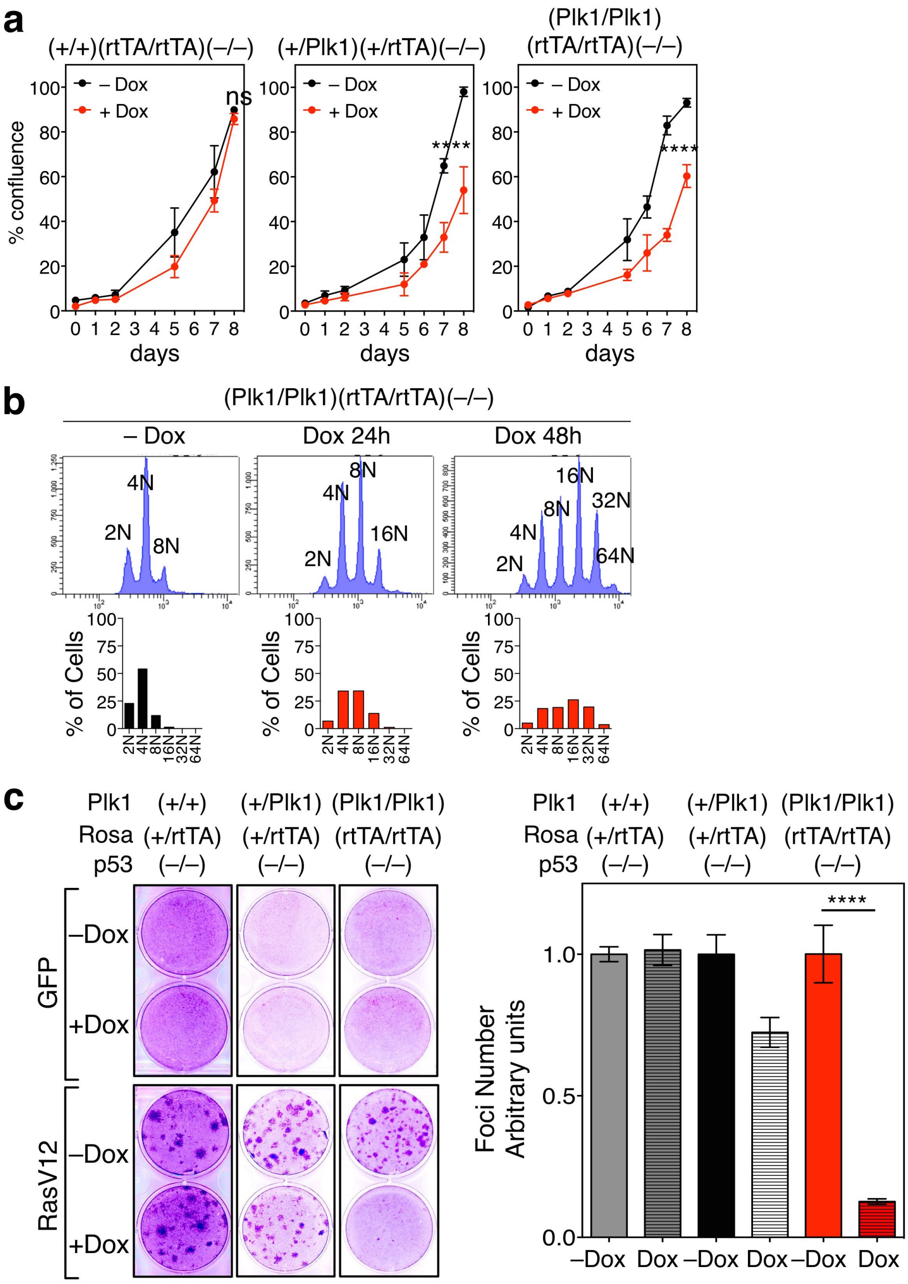


**Supplementary Fig. 3**. Inducible expression of Plk1 in p53–/– MEFs. **a** Quantification of confluence in control, (+/Plk1);(+/rtTA); p53(–/–) and (Plk1/Plk1);(rtTA/rtTA) p53(–/–)MEFs in the presence or absence of Dox during 8 days. ns, not significant; ****p<0.0001; Two-way ANOVA. **b** DNA content of (Plk1/Plk1); (rtTA/rtTA); p53(–/–) MEFs in the absence (–Dox) or the presence of Dox during 1 or 2 days. The percentage of cells with different ploidy levels is indicated in the histograms. **c** Focus formation assay of cells with the indicated genotypes after transfection with a GFP-expressing control vector or the HrasV12 oncogene. The number of foci normalized versus the control cultures is shown in the histogram. ****, p<0.0001; One-way ANOVA.

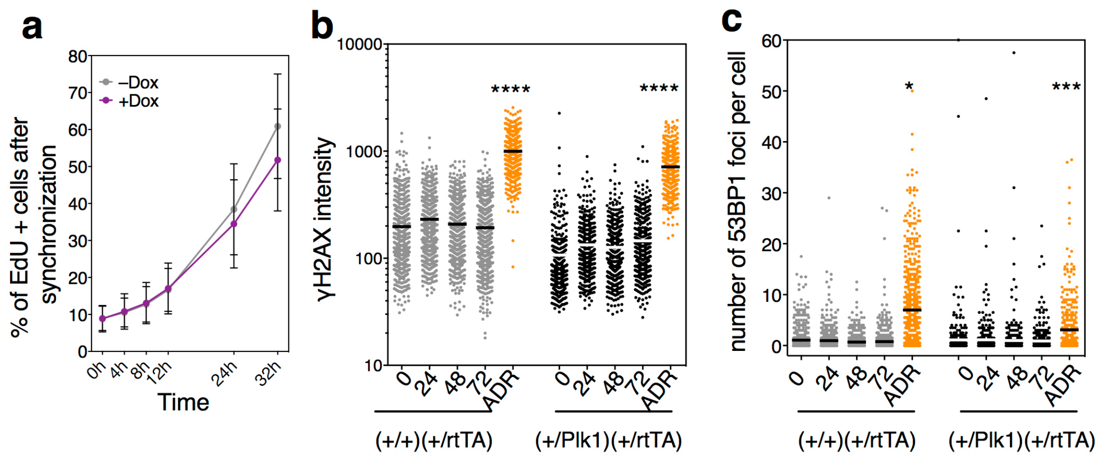


**Supplementary Fig. 4**. Plk1 induction does not induce DNA replication stress. **a** (+/Plk1);(+/rtTA) MEFs were grown up to confluency, and serum starved for 48h to stop the cell cycle at the G0 stage. Dox was added during the last 12h of serum starvation. Cells were then split and seeded in full serum media to force them to enter in the cell cycle, and analyzed for EdU incorporation by FACs, at the indicated times after cell cycle entry. Differences are not significant; 2-way ANOVA (n=4 replicates). **b,c** MEFs with the indicated genotypes were treated with Dox for the indicated h, and immunostained for γH2AX, 53BP1 and DAPI. Fluorescent mean intensity signal of γH2AX (**b**), and the number of 53BP1 foci per cell (**c**), was quantified with the OPERA high throughput microscope. Each dot in the columns represents an individual cell. Adriamycin (ADR) was added to cells as positive inductor of DNA damage. In **b,c** *, $p<0.5$; ***, $p<0.001$; ****, $p<0.0001$; One-way ANOVA.

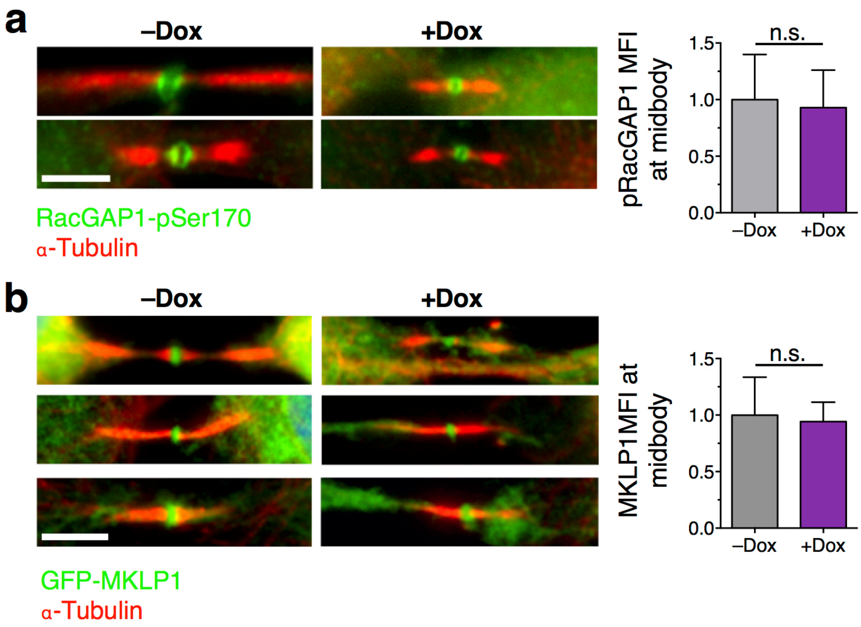


**Supplementary Fig. 5**. Plk1 induction does not disturb RacGAP1 activity or MKLP1 localization at the midbody. **a** (+/Plk1);(rtTA/rtTA) MEFs were untreated (–Dox) or treated for 24 h with Dox (+Dox). Cells were then immunostained for α-tubulin (red) and phospho-Ser170 RacGAP1 (green). Phospho-RacGAP1 mean fluorescence intensity at the midbody was quantified under the microscope. Data in –Dox sample were normalized to 1. (–Dox, n=11 cells; +Dox, n=34 cells). Scale bar, 5μm. **b** (+/Plk1);(rtTA/rtTA) MEFs were transfected with GFP-MKLP1 and treated for 24 h with Dox (+Dox) or left untreated (–Dox). Cells were then immunostained for α-tubulin (red), and GFP-MKLP1 (green) signal mean fluorescence intensity at the midbody was quantified under the microscope. Data in –Dox sample were normalized to 1. (–Dox, n=23 cells; +Dox, n=29 cells). In **a**,**b**, n.s., not significant, Student's t-test. Scale bar, 5μm.

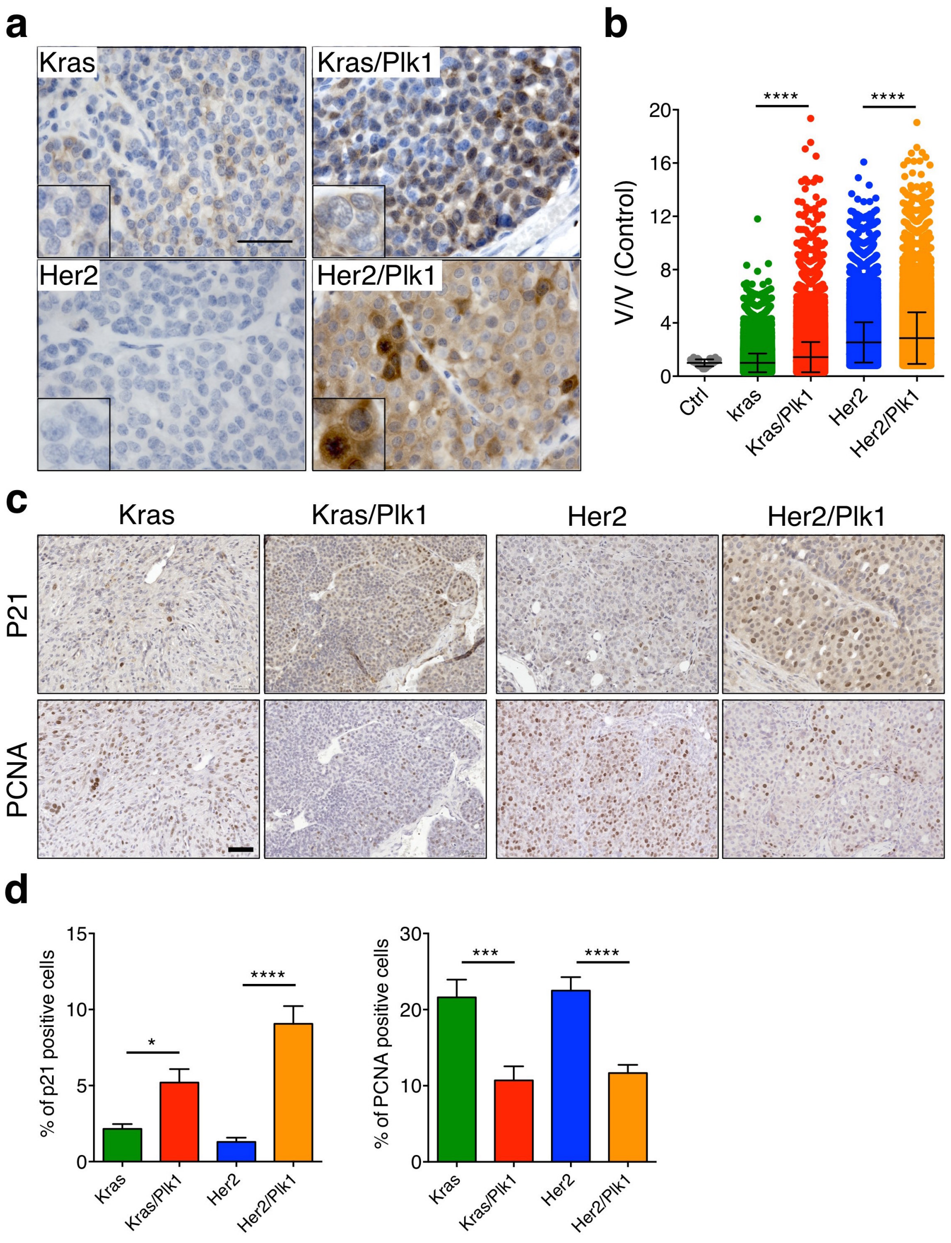


**Supplementary Fig. 6.** Phenotypic analysis of Plk1 expressing mammary tumors. **a** Immunohistochemistry against Plk1 on paraffin sections of mammary tumors. Scale bar, 20µm. **b** Quantification of nuclear volume of mammary tumor cells relative to control cells at 100 days on Dox (Ctrl, n=60; Kras, n=42846; Kras/Plk1, n=52359; Her2, n=31419; Her2/Plk1, n=28283 cells). Points represent individual nuclei. ****p < 0.0001, Kruskal-Wallis test, Dunn's multiple comparisons test. **c** Immunohistochemistry against p21 and PCNA in paraffin sections of mammary tumors in the indicated genotypes. **d** Quantification of the percentage of p21 positive cells per field of view (left graph), ordinary one-way ANOVA. *p < 0.05, ****p < 0.0001 and quantification of the percentage of PCNA positive cells per field of view (FOV) (right graph), ordinary one-way ANOVA. ***p < 0.001, ****p < 0.0001. (FOV analyzed minimum 30); points represent percentage of positive cells per FOV; 4 tumors analyzed per genotype.

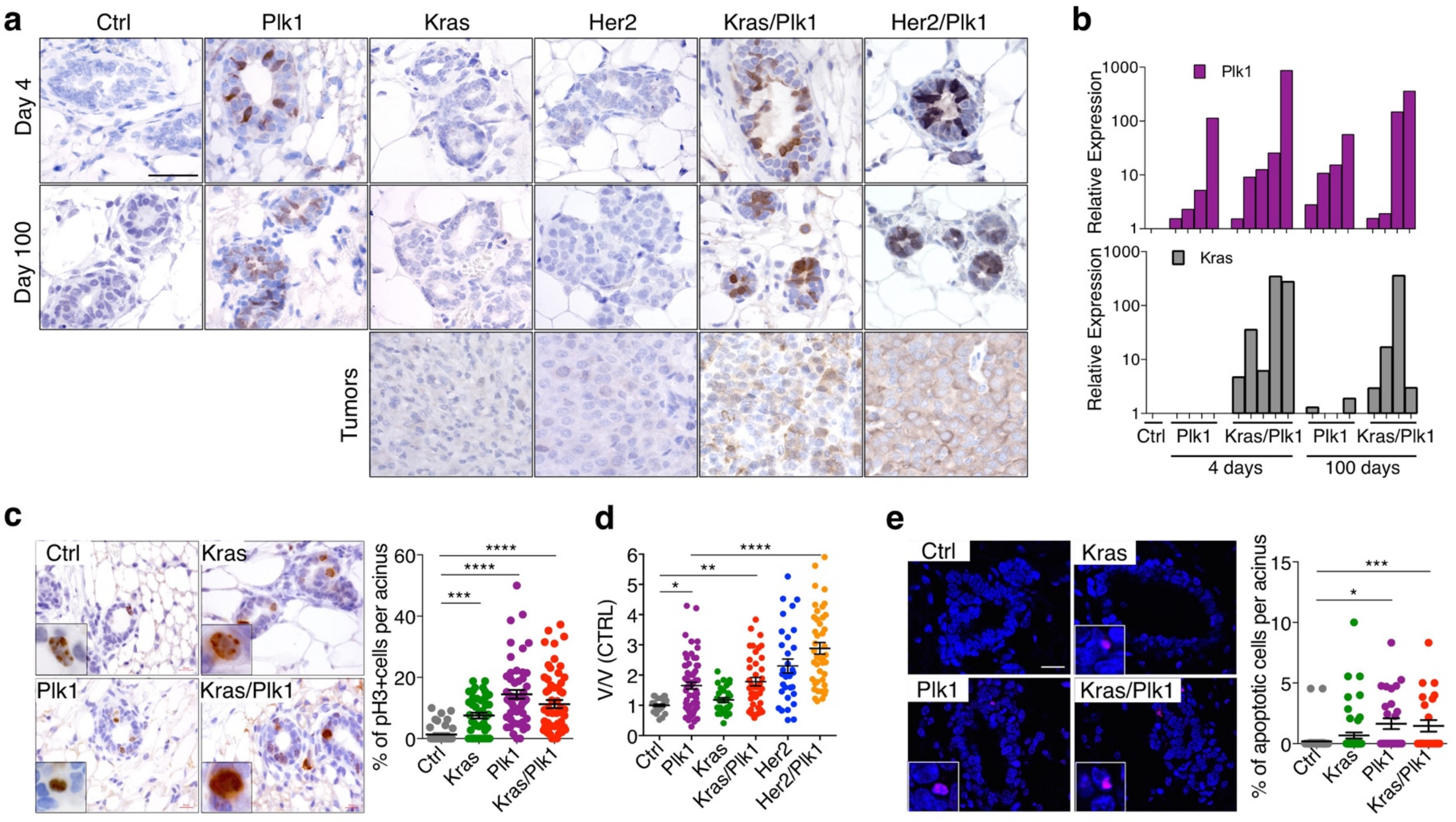


**Supplementary Fig. 7.** Cellular phenotypes associated to Plk1 overexpression in Kras-Her2 transgenic mice. **a** Immunohistochemistry against Plk1 in paraffin sections of mammary glands 4 days and 100 days on doxycycline and in the developed tumors. Scale bar, 50 µm. **b** Quantification by RT-qPCR showing relative expression of transgenic Plk1 and Kras$^{G12D}$ transcripts in mammary glands 4 days and 100 days on doxycycline. **c** Immunohistochemistry against phospho-H3 in paraffin sections of mammary glands 4 days on doxycycline. The histogram shows the percentage of pH3 positive cells per acinus (Ctrl, n=42 acini; Kras, n=42 acini; Plk1, n=53 acini; Kras/Plk1, n=58 acini). Each point represents single acini and total quantification includes minimum 6 animals per genotype. ***, p<0.001, ****, p<0.0001, Kruskal-Wallis test, Dunn's multiple comparisons test. **d** Quantification of nuclear volume relative to control cells at 4 days on doxycycline (Ctrl, n=30; Plk1, n=62 cells, Kras=30, Kras/Plk1=40, Her2=31, Her2/Plk1=43). Points represent single nuclear measurements. *, p<0.05; **, p<0.01; ****, p<0.0001 Kruskal-Wallis test, Dunn's multiple comparisons test. **e** Immunofluorescence using TUNEL kit in paraffin sections of mammary glands 4 days on doxycycline. The plot shows the quantification of percentage of TUNEL positive cells per acinus (Ctrl, n=51 acini; Kras, n=56 acini; Plk1, n=28 acini; Kras/Plk1, n=24 acini). Each point represents single acini and total quantification includes minimum 4 animals per genotype. *, p<0.05, ***, p<0.001, Kruskal-Wallis test, Dunn's multiple comparisons test.

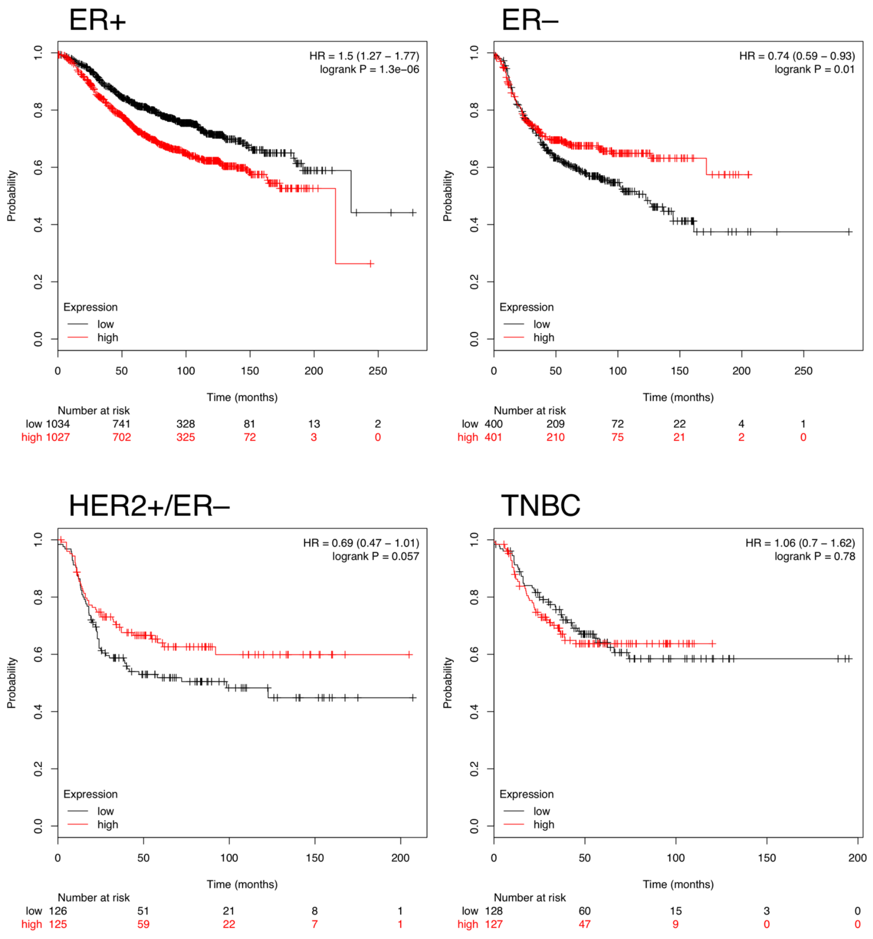


**Supplementary Fig. 8**. Differential prognostic value of Plk1 expression in mammary gland tumors. Plots represent the survival of patients with low (black) or high (red) tumoral levels of *PLK1* transcripts. Data were taken from Györffy et al. [1] and were analyzed using KMPlot (www.kmplot.com). ER+, estrogen receptor-positive (n=2061 patients); ER-, estrogen receptor-negative (n=801); HER2+/ER-, HER2-positive, ER-, estrogen receptor-negative (n=251); TNBC, triple negative breast cancer (n=255).

### Supplementary References

[1] Györffy, B., Lanczky, A., Eklund, A.C., Denkert, C., Budczies, J., Li, Q. & Szallasi, Z. An online survival analysis tool to rapidly assess the effect of 22,277 genes on breast cancer prognosis using microarray data of 1,809 patients. *Breast Cancer Res Treat*. **123**, 725-731. (2010)
